# Demographic changes and behavioural responses shape vulnerability to infectious disease outbreaks

**DOI:** 10.64898/2026.05.11.724461

**Authors:** Abbie Evans, William S. Hart, Eunok Jung, Kyeongah Nah, Karla Bonic-Babic, Sung-mok Jung, Robin N. Thompson

## Abstract

Demographic shifts are reshaping population age structures worldwide, with implications for infectious disease dynamics. Since contact patterns, susceptibility and infectiousness often vary by age, the risk that pathogen introductions initiate a substantial outbreak depends on the population’s age distribution and associated behavioural characteristics. We develop an age-structured mathematical model to estimate the risk that a single pathogen introduction leads to sustained transmission (the probability of a major outbreak) under long-term demographic transitions, incorporating changes in age-specific contact patterns and behavioural adaptation (specifically, extended workforce participation in an ageing society). Using the Republic of Korea (projected to become the world’s oldest population by 2050) as a case study, we show that population ageing may reduce the probability of a major outbreak due to older individuals’ lower contact rates. However, this effect is attenuated for pathogens with increasing susceptibility or infectiousness with age, and by factors such as later retirement that increase contacts among older individuals. These findings demonstrate that, while outbreak risks are affected by demographic changes, they are further modified by associated behavioural responses, highlighting the importance of accounting for demographic and socio-behavioural context when assessing future infectious disease outbreak risks.

**Author Summary:** In the early stages of an infectious disease outbreak, the risk that initial cases lead to substantial transmission is shaped by a range of factors including the characteristics of the host population. Demographic changes, such as population ageing, are transforming societies worldwide, yet their implications for infectious disease emergence remain unclear. Here, we show that population ageing may reduce the likelihood that imported infections trigger major infectious disease outbreaks due to lower contact rates among older individuals compared to younger individuals. However, this effect depends on how susceptibility, infectiousness and host behaviour vary with age. For example, if future older adults have increased social connectedness (e.g. due to a higher retirement age), then decreases in the outbreak risk may be offset. These findings underscore the need to account for demographic and socio-behavioural factors, in addition to biological factors, when assessing future outbreak risks and designing robust public health strategies, particularly in societies undergoing rapid demographic change.

## 1 Introduction

Population ageing is a defining demographic trend in many countries in the 21st century [1]. Falling fertility rates and increasing life expectancy have led to a rise in the median age in most countries, with a growing proportion of older adults and a shrinking proportion of younger individuals [2]. These demographic changes have wide-ranging implications for public health, including as a result of altered infectious disease dynamics. Transmission of many pathogens (e.g. SARS-CoV-2, influenza virus and respiratory syncytial virus) is driven by interactions between individuals. Since contact patterns vary with age [3], shifts in population age structures affect transmission dynamics. In addition, the impact of demographic change is further modulated by behavioural responses to varying population structures, as well as by age-specific biological characteristics that differ between pathogens (such as host susceptibility and infectiousness).

Mathematical models provide a framework for analysing how demographic and epidemiological processes interact. Previous epidemiological modelling studies have incorporated demographic context, demonstrating that changes in age structure can influence pathogen transmission dynamics and the resulting disease burden of outbreaks. For example, Møgelmose *et al*. [4] used an individual-based model to investigate the influence of demographic change on transmission of SARS-CoV-2 and influenza virus in Belgium, demonstrating that ageing populations may experience smaller outbreaks that cause larger numbers of deaths. Similarly, Geard *et al*. [5] analysed the Australian population from 1910 to 2010, showing that demographic transitions modify infection dynamics and the impacts of vaccination programmes.

Beyond disease burden, recent work has explored how population structures influence the risk that initial cases of disease in a host population lead to sustained local transmission, as opposed to an outbreak that fades out with few cases (Figure 1A). The “probability of a major outbreak” [6, 7, 8] has been estimated to quantify this risk, accounting for various types of population structure, including host populations stratified according to individuals’ immune status [9, 10] or spatial location [11]. In the context of age-structured populations, Nishiura *et al*. [12] demonstrated that the probability of a major outbreak differs depending on whether the index case is a child or adult, reflecting different social activity levels between these population groups. Extending this, Lovell-Read *et al*. [13] presented a framework for quantifying the probability of a major outbreak in age-structured populations in the context of non-pharmaceutical interventions against COVID-19 in the UK. Their work highlighted the importance of differences in contact rates and infectivity patterns between age groups when assessing outbreak risks, and demonstrated that the effectiveness of interventions depends on the age groups that are targeted.

**Figure 1:**
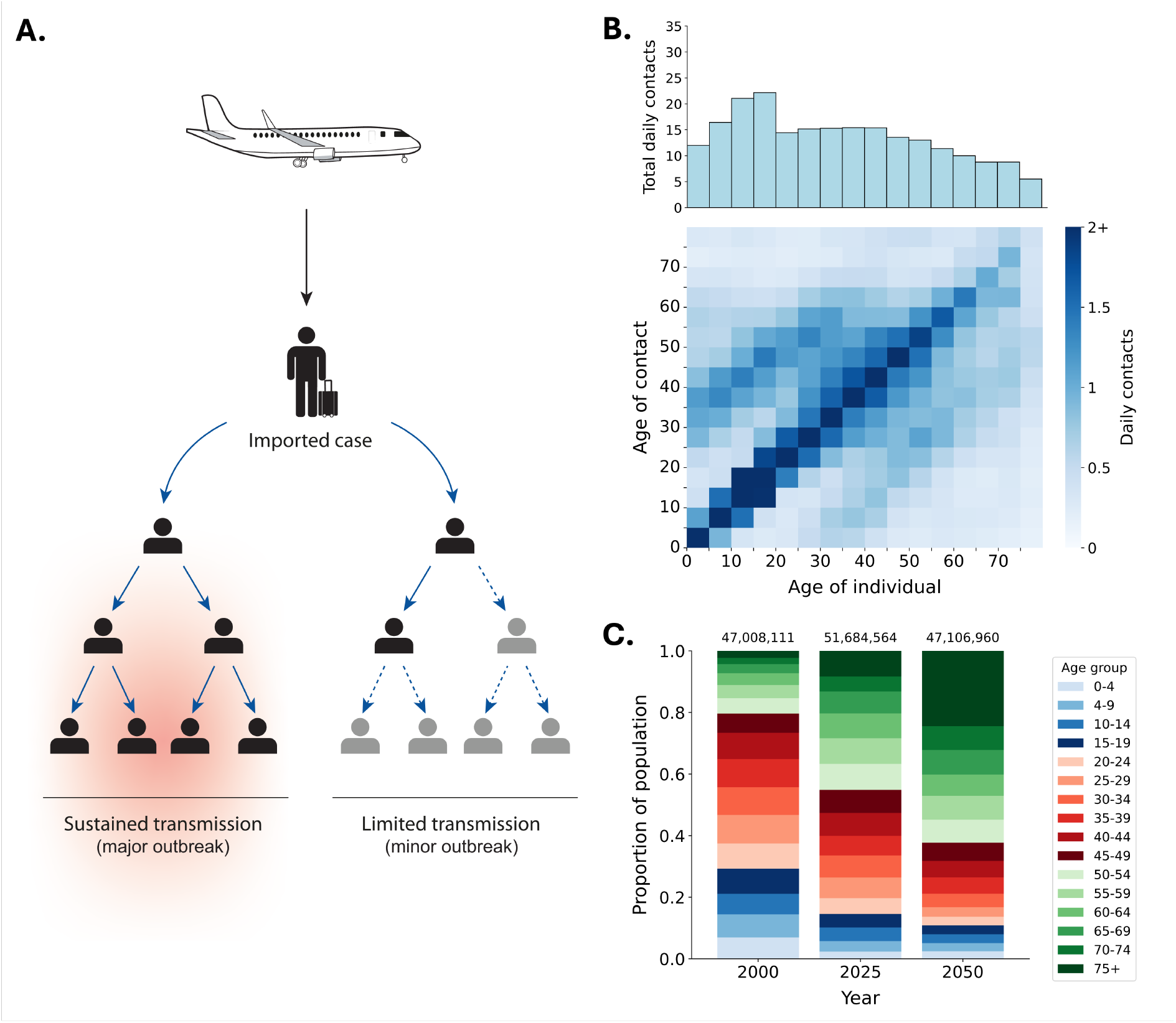
A. Schematic illustrating that, when an index case arrives in a host population, there are two possibilities for the outbreak that follows: either sustained transmission occurs (a major outbreak) or the outbreak fades out with few cases (a minor outbreak). B. The contact matrix in South Korea in 2020 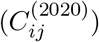, representing the expected daily number of unique contacts that an individual in the age group shown on the x-axis has with individuals in the age group shown on the y-axis. The graph shown on top represents the total expected daily number of unique contacts that an individual in the age group shown on the x-axis has (i.e. the column sum of the contact matrix). C. Age distribution of the population of South Korea in 2000, 2025 and 2050, based on United Nations estimates in 2000 and 2025 and projections for 2050 [18]. The values at the top of this panel indicate the estimated total population sizes in those years.

Despite increasing recognition that population structures influence outbreak risks, the effects of long-term demographic transitions and associated behavioural adaptations on these risks remain poorly understood. To address this gap, we develop an age-structured branching process transmission model and investigate how shifts in population age structure affect the probability of a major outbreak, considering pathogens with different age-specific patterns of susceptibility and infectiousness but comparable current transmission potential. As a case study, we focus on demographic changes in the Republic of Korea (hereafter, “South Korea”), which is one of the fastest ageing societies globally with fertility well below replacement and life expectancy continuing to rise [14, 15]. Our aim is not to generate precise country-specific projections of the probability of a major outbreak, but rather to use South Korea as an illustrative example to explore how demographic shifts associated with population ageing may qualitatively affect the probability of a major outbreak going forwards. We compare projections of the probability of a major outbreak in South Korea with analogous projections in the context of Nigeria; the latter was chosen as an illustrative example of a contrasting location where the population is generally younger and the current age structure is expected to remain similar in future. We further consider a scenario involving a behavioural response to population ageing; specifically, the extension of workforce participation to older adults. This reflects the possibility that individuals may live longer, healthier lives and remain socially and economically active. Through this approach, we explore how the interplay between population ageing and socio-behavioural adaptation affects the probability of a major outbreak, providing insights into future outbreak risks in a changing demographic landscape.

## 2 Methods

### 2.1 Mathematical model

We consider a population stratified into five-year age groups, with individuals aged 75 and over combined into a single age group. Hence, there are 16 age groups in total, made up of individuals aged 0-4, 5-9, and so on, up to 70-74, and 75 and over. Based on contact patterns in the year *y*, the expected number of infections in age group *j* generated by a single infected individual in age group *i* is 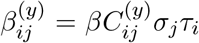, where 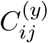 is the expected daily number of contacts that a single individual in age group *i* has with individuals in age group *j* (i.e. the relevant entry of the contact matrix in year *y*), *σ*_*j*_ reflects the susceptibility to infection of susceptible individuals in age group *j* and *τ*_*i*_ reflects the infectiousness of infected individuals in age group *i*. The basic reproduction number in the year *y*, 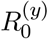, is the largest eigenvalue of the next generation matrix that has entries 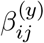 [16]. The parameter *β* (which is a composite parameter reflecting factors such as the probability of transmission per contact and the expected infectious period duration) determines the overall transmissibility of the pathogen, and is chosen to set a specified value of 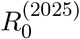 (see below).

To calculate the probability of a major outbreak in the year *y*, we follow the probability generating function method described by Southall *et al*. [6]. Specifically, we derive a system of non-linear simultaneous equations from which the probability of a major outbreak starting from a single infected individual in any given age group can be calculated, as explained below.

We first denote the probability of a major outbreak not occurring starting from a single index case in age group *i* in year *y* by 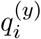. We assume that the total number of infections generated by an infected individual in age group *i* (which we denote *Z*_*i*_) follows a negative binomial distribution with mean 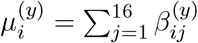 and dispersion parameter *k*. These infections are then distributed between age groups according to a multinomial distribution, where the probability that a specific infection is in age group *j* is 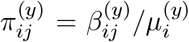. For a randomly chosen infection generated by an index case in age group *i*, the probability that a major outbreak does not follow from that infection is 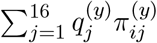 (which follows from conditioning on the possible age groups of that infection).

Hence, starting from a single infected individual in age group *i*, and conditioning on the number of infections generated by that individual,

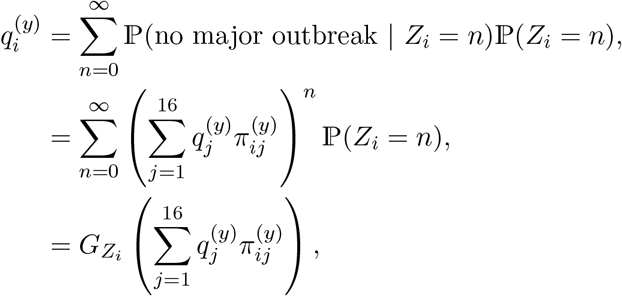

where 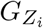 is the probability generating function of *Z*_*i*_.

Noting that the probability generating function of a negative binomial distribution with mean 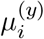 and dispersion parameter *k* is

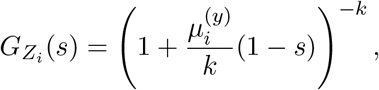

then, following some simple algebra,

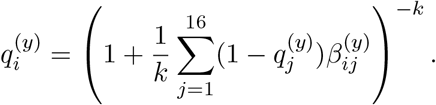

This equation holds for *i* = 1, 2, …, 16, giving a system of 16 equations that can be solved simultaneously using a numerical solver (in our analyses, we use the `fsolve` method from the `scipy.optimize` library in Python). The probability that a major outbreak does not occur starting from a single infected individual in age group *i* is the minimum non-negative solution for 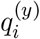 [6], and the probability of a major outbreak starting from a single infected individual in age group *i* is then 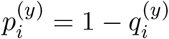.

We consider two metrics to quantify the overall probability of a major outbreak (over a range of possible index cases). First, we calculate the population average probability of a major outbreak,

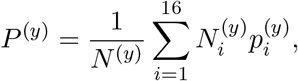

where *N* ^(*y*)^ is the total population size in year *y* and 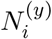 is the population size in age group *i* in year *y*. This represents the probability of a major outbreak starting from a single index case under the assumption that the probability that the index case is in age group *i* is equal to the proportion of the host population in that age group.

Second, we consider the contact-weighted average probability of a major outbreak,

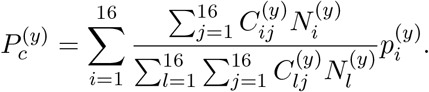

This quantity represents the probability of a major outbreak starting from a single infected individual, assuming that the probability that the index case is in age group *i* is proportional to the total expected number of contacts that all individuals in age group *i* have (i.e. summed across all individuals in age group *i*). This reflects the possibility that individuals in socially active age groups may be more likely to import the pathogen into the population (e.g. through travel or exposure to external infection sources).

In all of our analyses, except where stated, we assume that *k* = 1 so that the number of infections generated by a single infected individual follows a geometric distribution. However, we also investigate the robustness of our results to this choice, including considering scenarios in which *k <* 1 (which is representative of substantial super-spreading [10, 17]).

### 2.2 Model parameterisation

#### 2.2.1 Demographic parameters

The number of individuals in age group *i* in year *y*, 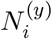, is chosen according to United Nations age demographic estimates and future projections for South Korea [18]. The expected daily number of contacts that an individual in age group *i* has with individuals in age group *j* in the year *y*, 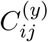, is estimated as follows. First, we consider the pre-COVID contact matrix for South Korea estimated by Prem *et al*. [3] with entries *C*_*ij*_. We then apply a pairwise correction to that contact matrix so that the expected total number of daily contacts that all individuals in age group *i* have with individuals in age group *j* is equal to the expected total number of daily contacts that all individuals in age group *j* have with individuals in age group *i*, as must be the case in reality [19]. We assume that the resulting contact matrix applied in 2020 (given that the source data [3] correspond to the beginning of 2020, we assume that the contact patterns had not yet been affected by the COVID-19 pandemic). We therefore set

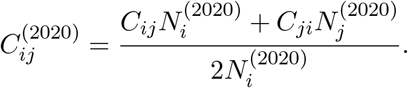

The resulting contact matrix for 2020, 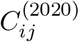, is shown in Figure 1B.

In order to evaluate the probability of a major outbreak in different years, we first estimate the contact matrix in year *y* (where we consider estimates for *y* = *{*2000, 2025, 2050*}*). To do this, we consider two alternative contact matrix projection methods (and we use the first of these methods throughout except where stated). The first method is the commonly applied density correction method [19, 20, 21, 22, 23, 24]. This method preserves assortativity (i.e. the relative preference of individuals in age group *i* for contacts with individuals in age group *j* compared to that expected under homogeneous mixing between individuals) and adjusts the contact matrix based on the ratio of the projected population density of individuals in the contact age group in year *y* and the density of individuals in the contact age group in the reference year (2020),

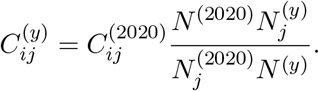

The second contact matrix projection method that we consider instead models a qualitatively different scenario in which individuals’ contact networks are assumed to remain similar going forwards. This reflects the possibility that, for example, school class sizes may remain similar even if the population age structure changes. Under that method, we assume that the contact matrix does not vary between years other than through a pairwise correction that we apply to ensure that, for the population age structure in year *y*, the expected total number of daily contacts that all individuals in any age group *i* have with individuals in any age group *j* matches the expected total number of daily contacts that all individuals in age group *j* have with individuals in age group *i*.

#### 2.2.2 Transmission parameters

For different pathogens, the age-dependence of individuals’ susceptibility to infection and their infectiousness once infected may differ. We therefore consider four different scenarios to capture a range of age profiles for these characteristics:

- **Uniform**. Under the uniform age profile, *σ*_*j*_ = 1 and *τ*_*j*_ = 1 for all age groups *j* = 1, 2, …, 16. In most of our analyses (except where stated), this is the baseline scenario for assessing the impacts of demographic and behavioural factors alone on the probability of a major outbreak.
- **Linearly increasing**. Susceptibility or infectiousness is assumed to increase linearly with age. Such a scenario is consistent with observations early in the COVID-19 pandemic that fewer cases were occurring among children than adults [25].
- **Linearly decreasing**. Susceptibility or infectiousness is assumed to decrease linearly with age, so that a disproportionate number of infections occur in younger age groups. This is representative of outbreak scenarios in which older cohorts have acquired partial immunity through prior exposure, as estimated during the 2009 influenza A (H1N1) pandemic [26].
- **U-shaped**. Both the youngest and oldest age groups are assumed to have elevated susceptibility or infectiousness, as seen for pathogens such as respiratory syncytial virus [27].

Under each sceario, we consider two subcases in which either susceptibility (*σ*_*j*_) or infectiousness (*τ*_*j*_) differs between age groups (*j*). To disentangle the effects of age-dependent susceptibility and age-dependent infectiousness, we do not consider scenarios in which *σ*_*j*_ and *τ*_*j*_ both vary with *j*.

The values of *σ*_*j*_ and *τ*_*j*_ used for each of these scenarios are given in Table S1. In each scenario, the value of the transmission parameter *β* is chosen so that 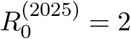 to ensure comparability across scenarios. In the different scenarios, the value of *β* is: *β* = 0.14 (uniform scenario), *β* = 0.15 (linearly increasing scenario), *β* = 0.11 (linearly decreasing scenario) and *β* = 0.15 (U-shaped scenario).

### 2.3 Behavioural responses to demographic changes

Under the density correction method, the intrinsic “preference” of an individual in age group *i* to have contacts with individuals in each age group does not vary through time. This naturally adjusts the expected number of contacts that an individual in a given age group is likely to have. For example, in South Korea, an individual in the 10-14 age group would be expected to have fewer contacts in future, since individuals in that age group tend to have most contacts with individuals of a similar age, and the proportion of individuals in the population in that age group will decline due to population ageing.

However, beyond these direct demographic effects, shifts in population age structure will likely drive associated behavioural responses that further modify future contact patterns. For example, societies with ageing populations are likely to require increased workforce participation among older adults (i.e. an increase in the retirement age), since the proportion of the population of working age will otherwise decrease. This could be aided by improvements in healthcare that enable individuals to live healthier lives for longer. The consequence of this socio-behavioural adaptation is that individuals in older age groups are likely to have more contacts compared to contact rates predicted by future changes in population age structure alone.

To analyse the effect of this possible behavioural response on the probability of a major outbreak, we model a scenario in which the retirement age increases from 60 in 2025 [28] to 75 in 2050. This adjustment is implemented in the contact matrix in 2050 by scaling entries corresponding to the expected daily number of contacts between an individual aged 60–74 and individuals aged 20–59, and entries corresponding to the expected daily number of contacts between an individual aged 20–59 and individuals aged 60–74. Specifically, these entries are scaled by multiplicative factor *ν*, yielding an increase in mixing between individuals aged 60–74 and individuals who are currently of working age. We set *ν* to be the ratio of the expected total daily number of contacts made by an individual aged 45–59 with individuals aged 20–59 in 2050 to the expected total daily number of contacts made by an individual aged 60–74 with individuals aged 20–59 in 2050 (before the retirement age adjustment is applied). This ensures that, after scaling, individuals aged 60–74 have similar numbers of contacts to those aged 45–59 in 2050.

### 2.4 Cross-country comparison

While our main focus in this article is demographic change in a country such as South Korea with an ageing population, we also generate analogous results under a contrasting demographic scenario. Specifically, we consider population age structure projections for Nigeria, which is a country with a younger population and an age structure that is projected to remain roughly stable in future. To analyse outbreak risks in 2000, 2025 and 2050 in that context, we apply the same methods as described above, but using United Nations age-specific population estimates and future projections for Nigeria [18], as well as the country-specific contact matrix for Nigeria in 2020 from the study by Prem *et al*. [3]. The impact of behavioural responses is not assessed in a Nigerian context, as its expected demographic stability (with a relatively high proportion of younger individuals) precludes the need to account for adaptations specific to an ageing society.

## 3 Results

### 3.1 Effect of population ageing on the probability of a major outbreak

To begin to investigate how outbreak risks are changing in South Korea as the population ages, we first generated contact matrices for 2000 (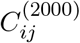; Figure 2A), 2025 (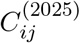; Figure 2B) and 2050 (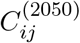; Figure 2C) using the density correction method. Under that method, by 2050, for all age groups, the expected number of contacts with individuals who are 75 and over is likely to increase, reflecting the shifting population age structure (Figure 1C).

**Figure 2:**
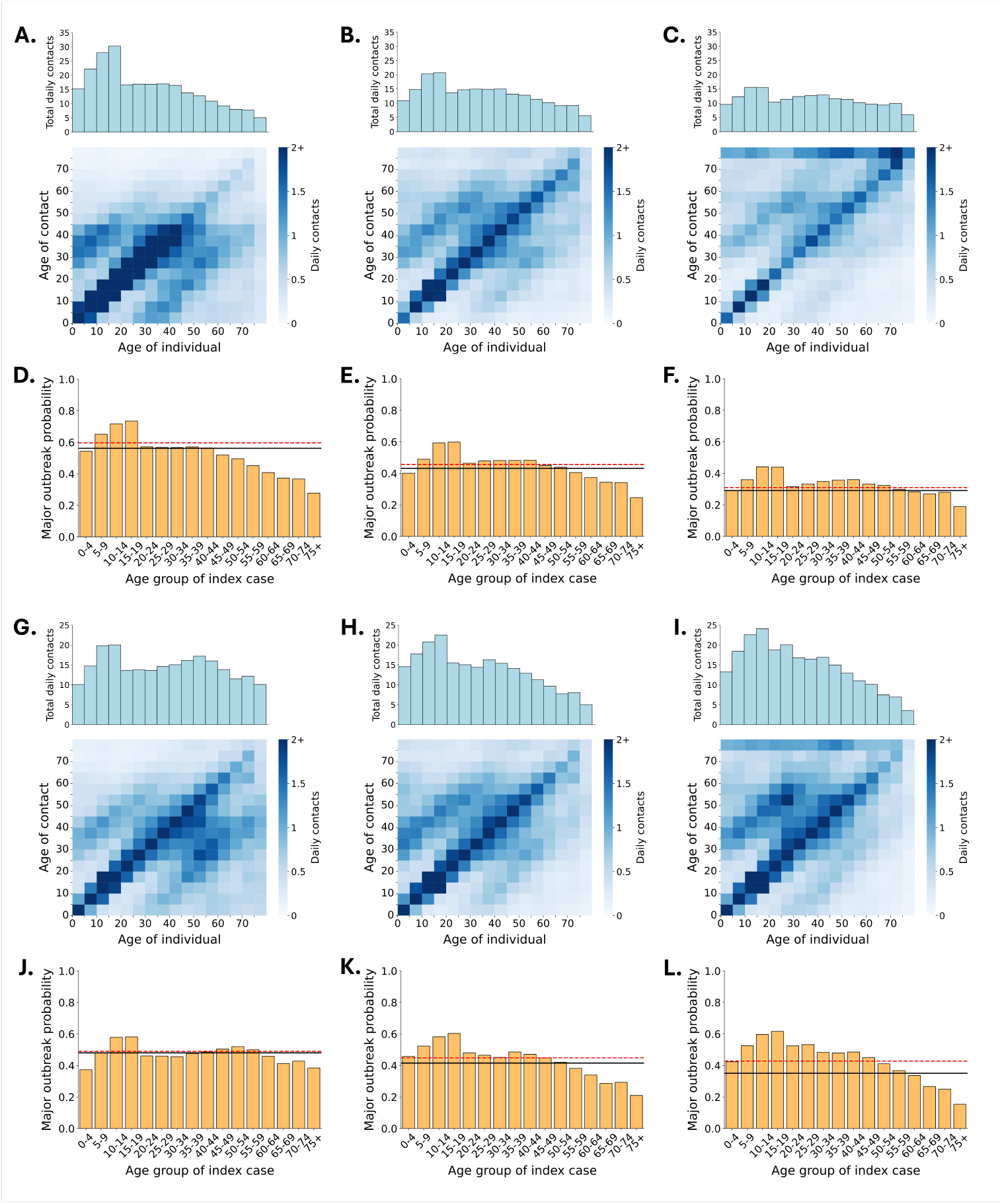
Changes in the probability of a major outbreak driven by demographic shifts in South Korea. A. The estimated contact matrix in South Korea in 2000 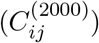, representing the expected daily number of unique contacts that an individual in the age group shown on the x-axis has with individuals in the age group shown on the y-axis. Here, we have used the density correction method. The graph shown on top represents the total expected daily number of unique contacts that an individual in the age group shown on the x-axis has (i.e. the column sum of the contact matrix). B. Analogous figure to panel A, but for 2025 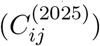. C. Analogous figure to panel A, but for 2050 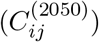. D. The probability of a major outbreak starting from a single infected individual in the specified age group (orange bars) in the year 2000. The population average probability of a major outbreak (black) and the contact-weighted average probability of a major outbreak (red dashed) are also shown. E. Analogous figure to panel D, but for 2025. F. Analogous figure to panel D, but for 2050. G–L. Identical to A–F, but using the pairwise correction contact matrix projection method.

Using these contact matrices, we calculated the probability of a major outbreak under our baseline scenario (uniform scenario; where both susceptibility and infectiousness are assumed to be independent of age), shown in Figure 2D–F. While the total population size is projected to be similar in 2050 compared to 2000 (Figure 1C), population ageing leads to a lower probability of a major outbreak, since older individuals tend to have fewer contacts than younger individuals. This conclusion held irrespective of the age of the index case; for any given age of the index case, the probability of a major outbreak always declined from 2000 to 2050 (Figure 2D–F, orange bars). Consequently, the population average probability of a major outbreak decreased over time (Figure 2D–F, black), as did the contact-weighted probability of a major outbreak (Figure 2D–F, red dashed).

We note that, across all three years considered (2000, 2025 and 2050), a major outbreak remains most likely to occur when the index case is in the 10-14 or 15-19 age groups. This can be attributed to the high contact rates of individuals in these age groups compared to individuals in other age groups (Figure 2A–C). However, the probability of a major outbreak for index cases in these age groups is projected to be substantially lower in 2050 than in previous decades. This decline is driven by age-based assortative mixing, whereby individuals tend to have a substantial proportion of their contacts with individuals of similar ages to themselves. Since the proportion of the population in the 10-14 and 15-19 age groups is expected to decrease by 2050, this not only means that the index case is expected to have fewer contacts, but also that the contacts of the index case will (on average) have fewer contacts. This compounding effect effectively thins the local transmission network of an index case aged 10-14 or 15-19, reducing the probability of a major outbreak.

We also conducted a supplementary analysis in which we analysed how the specified values of 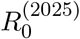 and *k* alter the population average probability of a major outbreak in 2000, 2025 and 2050 under our baseline model assumptions (Figure S1). Irrespective of the values of 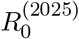 and *k*, under those assumptions we found that the population average probability of a major outbreak was projected to decrease in future.

We also conducted an additional analysis in which we made a qualitatively different assumption about how the contact matrix varies between years. Specifically, as described in Methods, we considered a scenario in which the contact matrix is assumed to remain relatively constant between years (due to the possibility that individuals’ contact networks could in principle remain similar even if the population is subject to demographic change). In that scenario, as expected, differences in the probability of a major outbreak between years are smaller than for the density correction method (Figure 2G–L), although the probability of a major outbreak nonetheless decreases in future (see Methods). Hereafter, and as before, we consider results from the density correction method.

### 3.2 Effect of age-based susceptibility or infectiousness on the probability of a major outbreak

For a range of pathogens, individuals’ susceptibility to infection is age-dependent. We investigated the impacts of different age profiles of susceptibility on the probability of a major outbreak, considering age profiles that were: (i) uniform (i.e. no variation in susceptibility with age); (ii) linearly increasing; (iii) linearly decreasing; and (iv) U-shaped. The relative susceptibility values for individuals in different age groups under each of these scenarios are shown in Figure 3A (and stated in Table S1).

**Figure 3:**
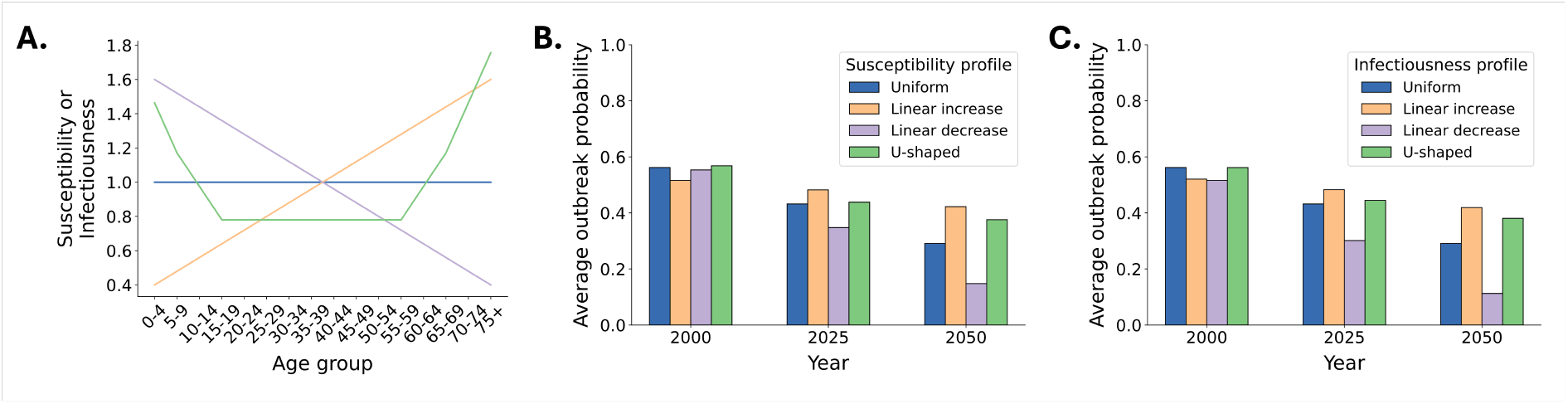
Temporal variation in the probability of a major outbreak for pathogens with different age-specific susceptibility and infectiousness profiles. A. Relative age-dependent susceptibility (*σ*_*j*_) or infectiousness (*τ*_*j*_) values considered in our analyses: uniform (blue), linearly increasing (orange), linearly decreasing (purple) and U-shaped (green). B. The population average probability of a major outbreak in different years and for different susceptibility profiles (with relative susceptibility values shown in panel A). C. The population average probability of a major outbreak in different years and for different infectiousness profiles (with relative infectiousness values shown in panel A).

The population average probability of a major outbreak in 2000, 2025 and 2050 for the four different susceptibility profiles is shown in Figure 3B, with 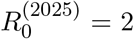 in each case. We found that the decrease in the population average probability of a major outbreak from 2000 to 2050 is largest for pathogens with susceptibility profiles that decrease with age (purple bars in Figure 3B). This is because, as the proportion of older individuals in the population increases, a greater proportion of all contacts in the population are then with less susceptible individuals and so are less likely to result in transmission.

In contrast, when we instead assumed that individuals’ susceptibility increases with age, the population average probability of a major outbreak only decreased slightly from 2000 to 2050 (orange bars in Figure 3B). This is because, by 2050, while individuals will typically have fewer contacts (due to an ageing population), more of these contacts will be with individuals with higher susceptibility. We note that the population average probability of a major outbreak was lower in 2000 when the linearly increasing susceptibility profile was considered than for any of the other age-dependent susceptibility profiles. This is because we fixed 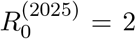 for all susceptibility profiles, and under the linearly increasing susceptibility profile, the relatively small proportion of older adults in 2000 (compared to in later years) translated to a larger number of contacts occurring with younger, less susceptible individuals, reducing the probability of a major outbreak.

In addition to considering different susceptibility profiles, we considered equivalent infectiousness profiles (then assuming that susceptibility was independent of age). As with the results for different susceptibility profiles, we found that the population average probability of a major outbreak decreased over time for all infectiousness profiles considered (Figure 3C), with the most substantial reduction in the population average probability of a major outbreak from 2000 to 2050 for the infectiousness profile that decreases with age.

We also considered the probability of a major outbreak for index cases of different ages under each of the susceptibility and infectiousness profiles (Figure S2). We found that there is often greater variation in the probability of a major outbreak between index cases of different ages, and between the different profiles, when infectiousness was assumed to vary with age than when susceptibility was assumed to vary with age. This is because the probability of a major outbreak is influenced substantially by the number of infections caused by the index case, and age-dependent infectiousness can lead to more variation in the number of infections generated by the index case between index cases of different ages. This arises because when infectiousness differs by age, it directly affects the expected number of secondary cases generated by the index case, potentially amplifying differences in the probability of a major outbreak between index cases of different ages. Age-dependent susceptibility, on the other hand, has a more diluted effect because contacts of the index case are distributed across multiple age groups with different susceptibilities.

### 3.3 Effect of behavioural responses on the probability of a major outbreak

Returning to the uniform susceptibility and infectiousness scenario, we next examined how changes in the retirement age (i.e. extended workforce participation in response to an ageing society) might influence the probability of a major outbreak. We found that an increased retirement age (to age 75) leads to a higher probability of a major outbreak in 2050 than when the retirement age remains at 60, particularly if the index case is between the ages of 60 and 74 (Figure 4). This increase is driven by increased workforce participation for those individuals, which creates a more connected contact network.

**Figure 4:**
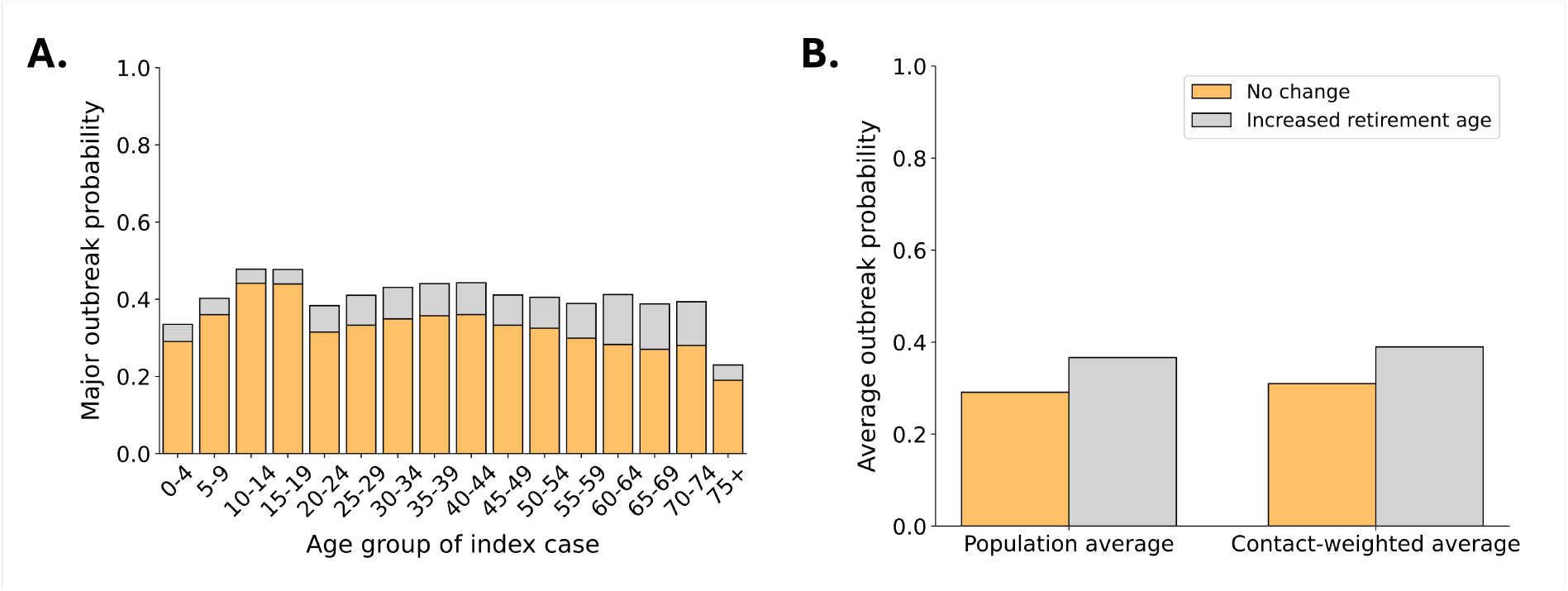
The effect of increasing the retirement age on the probability of a major outbreak. A. The probability of a major outbreak in 2050 without an increase in the retirement age (orange bars; as in Figure 2F). The probability of a major outbreak in 2050 when the retirement age is instead increased to 75 (grey bars). B. The population average probability of a major outbreak in 2050 and the contact-weighted probability of a major outbreak in 2050, either without (orange bars) or with (grey bars) an increase in the retirement age from 60 to 75.

However, even with the higher retirement age in 2050, we found that both the population average probability of a major outbreak and the contact-weighted probability of a major outbreak were lower in 2050 than in 2025. For example, the population average probability of a major outbreak was 0.43 in 2025 and 0.37 in 2050 when assuming that individuals retire at age 75.

### 3.4 Cross-country comparison

Although the probability of a major outbreak is expected to decrease in South Korea under the density correction method, this result depends on the projected population age structure. To compare the projected probability of a major outbreak in South Korea against the analogous quantity in a country with a contrasting demographic scenario, we considered the probability of a major outbreak in Nigeria, again setting 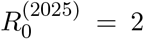. This comparison is intended to illustrate the effects of different projected population dynamics, rather than to provide a direct comparison between likely future outbreak risks in these two specific countries. Specifically, we do not account for differences beyond population age structure that might also affect outbreak risks, such as cultural or economic differences. In Nigeria, age structure is projected to remain relatively similar up to 2050, despite a substantial growth in the total population size (Figure 5A). Consistent with this, projections of age-specific contact patterns using the density correction method show minimal changes over time, reflecting the relatively stable proportion of individuals in each age group. As a result, we find little temporal variation in the probability of a major outbreak in Nigeria (Figure 5B–D). While the populations of most countries globally are expected to undergo population ageing by 2050, this result demonstrates that the resultant lower outbreak risk is not ubiquitous.

**Figure 5:**
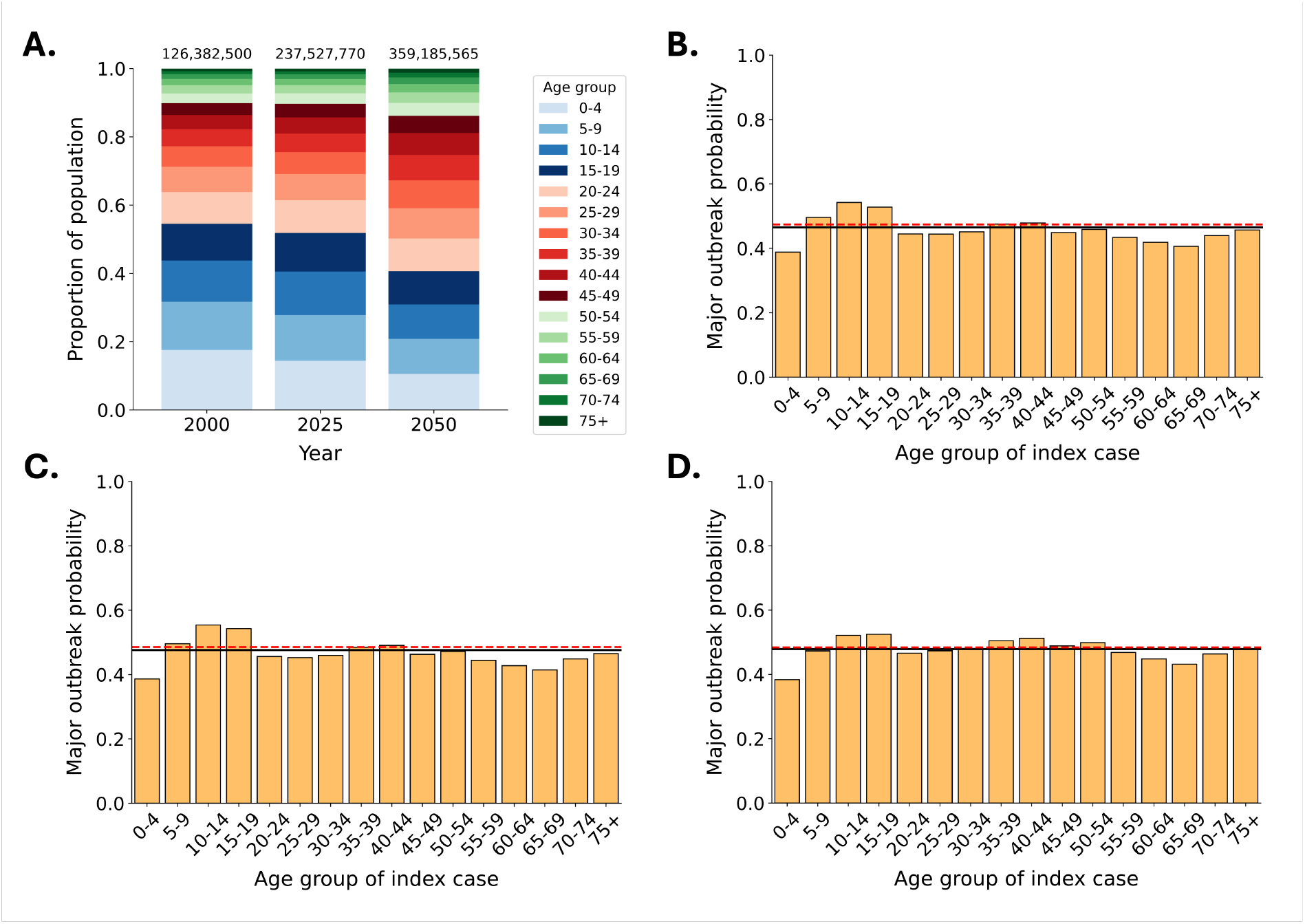
Changes in the probability of a major outbreak in Nigeria. A. Age distribution of the population of Nigeria in 2000, 2025 and 2050, based on United Nations estimates in 2000 and 2025 and projections for 2050 [18]. The values at the top of this panel indicate the estimated total population sizes in those years. B. The probability of a major outbreak starting from a single infected individual in the specified age group (orange bars) in the year 2000. The population average probability of a major outbreak (black) and the contact-weighted average probability of a major outbreak (red dashed) are also shown. C. Analogous figure to panel B, but for 2025. D. Analogous figure to panel B, but for 2050.

## 4 Discussion

Emerging and re-emerging pathogens pose a substantial threat to public health [29]. Understanding the circumstances in which pathogen introductions are likely to initiate sustained local transmission remains a fundamental challenge [30, 31, 32]. In this study, we used an age-structured branching process model to quantify how transmission risks are affected by the interactions between shifting demographics, pathogen characteristics and socio-behavioural adaptation in evolving societal landscapes.

We found that demographic changes alone can fundamentally alter outbreak risks. In demographic scenarios with populations similar to South Korea, and utilising a commonly used contact matrix projection method, the transition towards an older population is projected to reduce the probability of a major outbreak (Figure 2A–F). This decline is driven by the increasing proportion of older individuals (who have fewer contacts than younger individuals), which acts to thin the population-level transmission network. While different pathogens (represented by varying age-specific profiles of susceptibility and infectiousness) yield distinct results, underlining the importance of pathogen-specific transmission characteristics when estimating the future potential for outbreaks, a temporal decrease in the probability of a major outbreak was observed across all age-dependent susceptibility and infectiousness scenarios that we considered (Figure 3).

While population ageing could reduce the risk that outbreaks occur, our projections indicate that the extent of this decrease depends on how the socio-behavioural landscape evolves. Demographic shifts may necessitate structural adaptations, such as extending workforce participation to older individuals to mitigate labour shortages, that reconnect the transmission networks of older individuals. Increased activity of older individuals acts to increase the projected probability of a major outbreak (Figure 4), although in the case study of South Korea that we considered we found that the probability of a major outbreak in 2025 exceeded the projected probability of a major outbreak in 2050 even when an increase in the retirement age was accounted for. We note, however, that a range of other factors may affect future contact rates, with potentially contrasting effects. For example, higher workforce participation may not be required in future; increased automation (potentially facilitated by artificial intelligence) may reduce future labour demand and/or change working arrangements. Counteracting this, as populations age, a larger proportion of older individuals may reside in communal living settings, increasing contact rates among older individuals. Similar to the effect of increased workforce participation, these elevated contact rates could partially offset the projected potential reduction in the probability of a major outbreak associated with population ageing. Nonetheless, the increase in the probability of a major outbreak associated with a higher retirement age demonstrates the importance of incorporating socio-behavioural adaptation into epidemiological frameworks that are used to plan future disease surveillance strategies and outbreak responses.

The role of population ageing in shaping the future probability of a major outbreak in South Korea was particularly evident when we contrasted our main results against similar projections for Nigeria. Despite substantial projected growth in the total population size, Nigeria’s younger population with higher contact rates yielded relatively stable projections of the probability of a major outbreak going forwards (Figure 5). This highlights the importance of accounting for country-specific changes in population structure when assessing future outbreak risks.

It should be noted that a reduction in the probability of a major outbreak does not necessarily translate into lower disease burden if an outbreak occurs. If the risk of severe outcomes associated with infection increases with age, as observed during the COVID-19 pandemic [33], demographic transitions towards older populations may lead to less frequent and smaller, but potentially more severe, outbreaks (e.g. outbreaks with larger numbers of hospitalisations and deaths). This may require ageing societies to prioritise enhancing healthcare capacity to manage the high burden associated with outbreaks, as well as the development and deployment of vaccines and other pharmaceutical interventions to reduce the prevalence of severe infection outcomes. Consequently, from a public health policy perspective, potentially reduced outbreak risks due to ageing populations should not lead to complacency, as they may coincide with increased vulnerability to healthcare system strain when major outbreaks occur. We note that a similar result has previously been observed in the context of outbreaks with substantial transmission heterogeneity between individuals (super-spreading); this phenomenon leads to less frequent but more explosive outbreaks [17].

As with any modelling analysis, our study involved assumptions and simplifications. For example, we relied on projected contact matrices and assumed that these projections adequately represent future population mixing patterns. While the contact matrix projections underlying most of our results were based on a commonly used density correction method, contact networks may differ from these projections in future, motivating our additional analysis in which we assumed that contact matrices would instead remain similar in future (Figure 2G–L). Empirical evidence about precisely how changing population age structures affect contact patterns is currently limited, and longitudinal contact surveys conducted over sufficiently long periods to identify clear patterns are needed. However, it might be expected that current societal trends in South Korea – for example decreasing household sizes and increasing use of contactless services – act to amplify our finding that outbreak risks may decrease in future.

In addition, although we considered a range of age-dependent susceptibility and infectiousness profiles, some pathogens may exhibit more complex age-related effects. Nonetheless, the use of these different profiles enabled us to assess possible qualitative changes in the probability of a major outbreak in future for pathogens with different age-dependent transmission risks. Lastly, although our analysis focused on a scenario in which a single infected individual is introduced into the host population, providing us with a per-introduction estimate of the probability of a major outbreak, in reality multiple pathogen introductions may occur. The probability of a major outbreak can be calculated straightforwardly in the context of multiple pathogen introductions using the same approach as used here. For example, in year *y*, if the probability that a major outbreak will not occur starting from a single introduced infected individual in age group *i* is 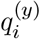, then the probability of a major outbreak starting from two infected individuals in age groups *i* and *j* is 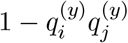, assuming independent transmission lineages.

Despite these limitations, our results provide insights into how outbreak risks are changing due to population ageing and associated behavioural adaptations, in addition to pathogen-specific characteristics. Together, these elements shape the future risk posed by emerging and re-emerging pathogens by influencing transmission networks. Future outbreak risk assessments should therefore adopt an integrated perspective that accounts for demographic and socio-behavioural drivers. Understanding these interactions is essential for designing robust public health strategies, particularly in societies undergoing rapid demographic change.

## Authors’ contributions

AE: methodology, formal analysis, investigation, visualisation, validation, writing – original draft, writing – review and editing.

WSH: methodology, supervision, writing – review and editing.

EJ: writing – review and editing.

KN: writing – review and editing.

KB-B: visualisation, writing – review and editing.

SJ: conceptualization, methodology, writing – review and editing.

RNT: conceptualization, methodology, supervision, writing – original draft, writing – review and editing.

## Data availability

The computing code used to perform the analyses in this article is available at https://github.com/abbieevans/korea-outbreak-risks. All code was written in Python v. 3.13.1.

## Acknowledgements

The authors are grateful to members of the Infectious Disease Modelling group in the Mathematical Institute in Oxford for helpful discussions throughout this project.

