## Supplementary Information for "Demographic changes and behavioural responses shape vulnerability to infectious disease outbreaks"

Abbie Evans, William S. Hart, Eunok Jung, Kyeongah Nah, Karla Bonic-Babic, Sung-mok Jung, Robin N. Thompson

#### 1 Supplementary Table

| **Age group ($j$)** | **Susceptibility ($\sigma_j$) or Infectiousness ($\tau_j$)** | | | |
|---|---|---|---|---|
| | Uniform scenario | Linearly increasing scenario | Linearly decreasing scenario | U-shaped scenario |
| 0–4 ($j = 1$) | 1.0 | 0.4 | 1.6 | 1.46 |
| 5–9 ($j = 2$) | 1.0 | 0.48 | 1.52 | 1.17 |
| 10–14 ($j = 3$) | 1.0 | 0.56 | 1.44 | 0.96 |
| 15–19 ($j = 4$) | 1.0 | 0.64 | 1.36 | 0.78 |
| 20–24 ($j = 5$) | 1.0 | 0.72 | 1.28 | 0.78 |
| 25–29 ($j = 6$) | 1.0 | 0.8 | 1.2 | 0.78 |
| 30–34 ($j = 7$) | 1.0 | 0.88 | 1.12 | 0.78 |
| 35–39 ($j = 8$) | 1.0 | 0.96 | 1.04 | 0.78 |
| 40–44 ($j = 9$) | 1.0 | 1.04 | 0.96 | 0.78 |
| 45–49 ($j = 10$) | 1.0 | 1.12 | 0.88 | 0.78 |
| 50–54 ($j = 11$) | 1.0 | 1.2 | 0.8 | 0.78 |
| 55–59 ($j = 12$) | 1.0 | 1.28 | 0.72 | 0.78 |
| 60–64 ($j = 13$) | 1.0 | 1.36 | 0.64 | 0.96 |
| 65–69 ($j = 14$) | 1.0 | 1.44 | 0.56 | 1.17 |
| 70–74 ($j = 15$) | 1.0 | 1.52 | 0.48 | 1.46 |
| 75+ ($j = 16$) | 1.0 | 1.6 | 0.4 | 1.76 |

Table S1: The relative susceptibility ($\sigma_j$) or infectiousness ($\tau_j$) values used in each of the scenarios described in the main text (see Figure 3 in the main text). Each scenario (except for the uniform scenario) consists of two subcases, with either $\sigma_j$ or $\tau_j$ varying between age groups ($j$).

### 2 Supplementary Figures

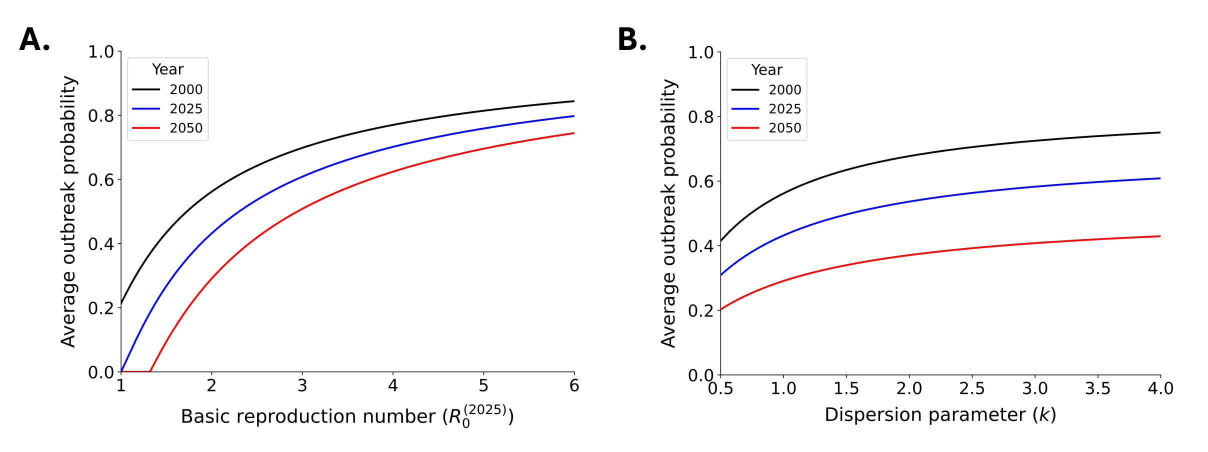


Figure S1: **Dependence of the population average probability of a major outbreak on values of $R_0^{(2025)}$ and $k$.** The population average probability of a major outbreak in 2000 (black), 2025 (blue) and 2050 (red) as a function of: A. The assumed value of the basic reproduction number in 2025 ($R_0^{(2025)}$). B. The assumed value of the dispersion parameter of the negative binomial offspring distribution ($k$). Here, as in most of our results, we applied the density correction method.

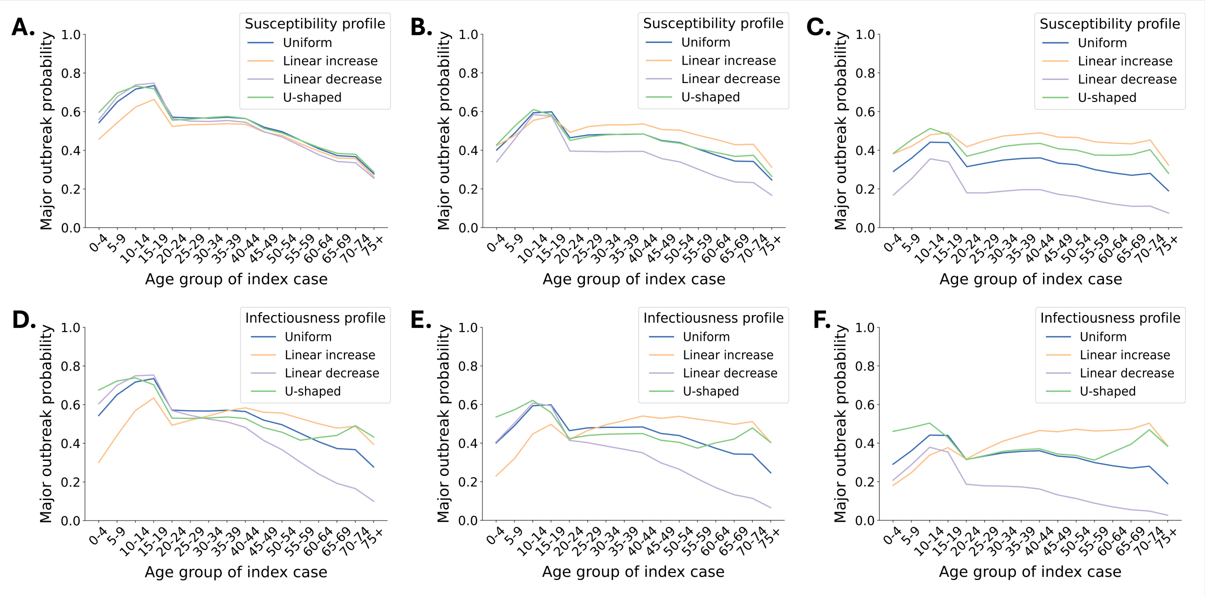


Figure S2: **Effects of different susceptibility and infectiousness profiles on the probability of a major outbreak.** A. The probability of a major outbreak in South Korea in 2000 starting from a single infected individual in the specified age group, for the uniform (blue), linearly increasing (orange), linearly decreasing (purple) and U–shaped (green) susceptibility profiles. B. Analogous figure to panel A, but for 2025. C. Analogous figure to panel A, but for 2050. D. Analogous figure to panel A, but instead considering different infectiousness profiles. E. Analogous figure to panel B, but instead considering different infectiousness profiles. F. Analogous figure to panel C, but instead considering different infectiousness profiles.
